# Overlooked roles of DNA damage and maternal age in generating human germline mutations

**DOI:** 10.1101/327098

**Authors:** Ziyue Gao, Priya Moorjani, Thomas Sasani, Brent Pedersen, Aaron Quinlan, Lynn Jorde, Guy Amster, Molly Przeworski

## Abstract

Although the textbook view is that most germline mutations arise from replication errors, when analyzing large *de novo* mutation datasets in humans, we find multiple lines of evidence that call that understanding into question. Notably, despite the drastic increase in the ratio of male to female germ cell divisions after the onset of spermatogenesis, even young fathers contribute three times more mutations than young mothers, and this ratio barely increases with parental ages. This surprising finding points to a substantial contribution of damage-induced mutations. Indeed, C to G transversions and CpG transitions, which together constitute one third of all mutations, show genomic distributions and sex-specific age dependencies indicative of doublestrand break repair and methylation-associated damage, respectively. Moreover, the data indicate that maternal age at conception influences the mutation rate both because of the accumulation of damage in oocytes and potentially through an influence on the number of postzygotic mutations.

## Introduction

Despite the fundamental importance of germline mutation as the source of heritable diseases and driver of evolution, its genesis remains poorly understood (Huttley *et al.*, 2000; Kumar and Subramanian, 2002) In principle, *de novo* point mutations could arise either directly from errors made while copying an intact DNA template (i.e., be “replication driven”) or from damage of the template or free nucleotides that occurred before DNA replication (be “damage-induced”) or an interaction of the two (e.g., Poulos, Olivier and Wong, 2017). Characterizing the relative importance of these mutation sources is of inherent interest and carries many implications, including for understanding the erratic behavior of the molecular clock used to date evolutionary events (Wu and Li, 1985; Hwang and Green, 2004; Wilson-Sayres *et al.*, 2011); for the nature of selection pressures on replication and repair machinery (Lynch, 2010; Lynch *et al.*, 2016); and in humans, for predicting recurrence risks of Mendelian diseases and disease burdens (Crow, 1997; Acuna-Hidalgo, Veltman and Hoischen, 2016).

Since germline mutation is extremely difficult to study directly, our understanding of the process is based on experiments in unicellular organisms or on the dependence of mutation rates in children on their parental ages in humans and other mammals (Ségurel, Wyman and Przeworski, 2014; Lindsay *et al.*, 2016; Lynch *et al.*, 2016; Harland *et al.*, 2017). Notably, the textbook view that replication errors are the primary source of human mutations (Drost and Lee, 1995; Li *et al.*, 1996; Crow, 2000; Strachan and Read, 2010) often invokes the increase in the number of germline mutations with paternal age (Crow, 2000; Drost & Lee, 1995). However, this increase can arise not only from DNA replication in spermatogonial stem cells, but also from other metabolic activities associated with cell division or from unrepaired damage that accrues with the passage of time (Müller, 1954). A further complication is that the rate at which unrepaired DNA lesions are converted into mutations is affected by DNA replication and thus can depend on the cell division rate (Gao *et al.*, 2016; Seplyarskiy *et al.*, 2017).

Insight into the genesis of mutations can also be gained by contrasting male and female mutation patterns, which reflect distinct development trajectories and epigenetic dynamics. To apply this approach, we re-analyzed *de novo* mutation (DNM) data from a recent study of over 1500 parent-offspring trios (Jónsson, Sulem, Kehr, *et al.*, 2017) and contrasted the properties of paternal and maternal mutations. We tested the hypothesis that germline mutations are primarily replicative in origin by asking: How well do male and female mutations track the numbers of germ cell divisions? Do the dependencies of the mutation rate on the parent’s sex and age differ by mutation types and if so, why? Is the higher male contribution to *de novo* mutations explained by replication errors in dividing male germ cells?

## Results

### The ratio of paternal to maternal mutations is already high for young parents and is stable with parental age

If mutations are replication-driven, the ratio of male to female germline mutations (also known as the male mutational bias, α) should reflect the ratio of the number of germ cell divisions in the two sexes. While male and female germlines are thought to undergo similar numbers of mitotic cell divisions by the onset of puberty (∼30-35 divisions), thereafter the ratio of male-to-female cell divisions increases rapidly with age, because of frequent mitotic divisions of spermatogonial stem cells (an estimated 23 divisions per year) and the absence of mitotic division of female germ cells over the same period (Vogel and Rathenberg, 1975; Drost and Lee, 1995). Thus, α is expected to increase substantially with parental age at conception of the child, even though it need not be strictly proportional to the cell division ratio, if per cell division mutation rates vary across developmental stages (Ségurel, Wyman and Przeworski, 2014; Gao *et al.*, 2016).

To test this prediction, we analyzed autosomal DNM data from 1548 Icelandic trios (Jónsson *et al.*, 2017; henceforth, the “deCODE dataset”), initially focusing on phased mutations, i.e., the subset of mutations for which the parental origin of the DNM had been determined by either transmission to third-generation individuals or linkage to variants at nearby heterozygous polymorphism in reads. Given that the phasing rate differs across trios (Fig. S1), we considered the fraction of paternal mutations in all phased DNMs and compared this fraction against the father’s age (see Materials and Methods). We found that, for trios with similar paternal age, G_P_, and maternal, G_M_ (where 0.9<G_P_/G_M_<1.1), as is the case for most families in the dataset, the paternal contribution to mutations is strikingly stable with paternal age (Fig. 1). The average contribution of paternal mutation lies around 75-80%, i.e., α = 3-4 across paternal ages, with a similar median α seen for children of parents aged 40 as aged 20 for example. This result remains after excluding C>G mutations, which were previously reported to increase disproportionately rapidly with maternal age (Jónsson, Sulem, Kehr, *et al.*, 2017)(Fig. S2). Moreover, the same result is seen in an independent DNM dataset containing 816 trios (Goldmann *et al.*, 2016; Wong *et al.*, 2016), which are also mostly of European ancestry (henceforth the “Inova dataset”; Fig. S3). The finding of a relatively stable α of 3-4 with parental ages calls into question the widespread belief that spermatogenesis drives the male bias in germline mutations (Crow, 1997; Makova and Li, 2002; Ségurel, Wyman and Przeworski, 2014).

**Figure 1.**
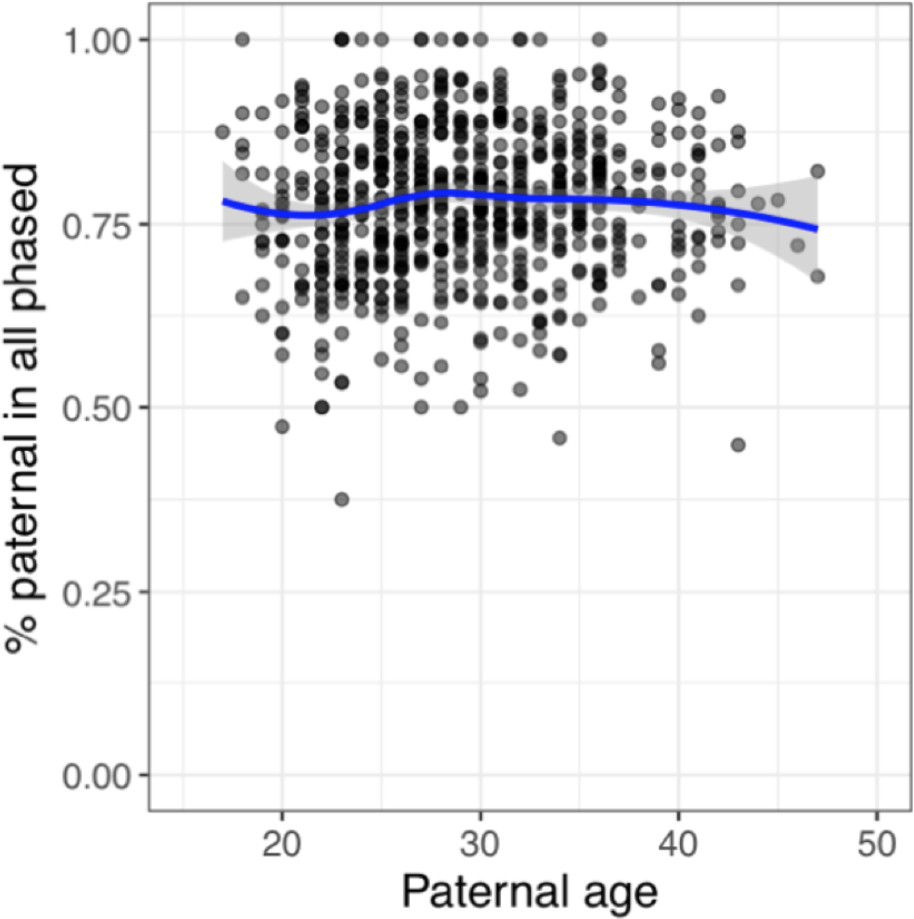
The fraction of paternal mutations among phased mutations, as a function of paternal age at conception. Each point represents the data for one child (proband) with at least three phased mutations and similar parental ages (paternal-to-maternal age ratio between 0.9 to 1.1; 719 trios total). The blue line is the LOESS curve fitted to the scatterplot, with the shaded area representing the 95% confidence interval (calculated with the geom_smooth function in R with default parameters).

A stable α implies that the number of maternal mutations increases with the mother’s age almost at the same relative rate as paternal mutations with father’s age. To obtain more precise estimates of parental age effects, we modeled paternal and maternal age effects jointly, leveraging information from both phased and unphased mutations in the deCODE dataset. Briefly, we modeled the expected number of mutations in a parent as a linear function of her (his) age at conception of the child, and assumed that the observed number of maternal (paternal) mutations follows a Poisson distribution. We further modeled the number of maternal (paternal) mutations that were successfully phased as a binomial sample of DNMs (see Materials and Methods). We then estimated the sex-specific yearly increases with parental ages by maximum likelihood (Table S1). Uncertainty in the estimates was evaluated by bootstrap resampling of trios. We confirmed earlier reports that the mutation counts on the paternal and maternal sets of chromosomes increased with father and mother’s ages, respectively (Fig. 2A)(Goldmann *et al.*, 2016; Wong *et al.*, 2016; Jónsson, Sulem, Kehr, *et al.*, 2017).

**Figure 2.**
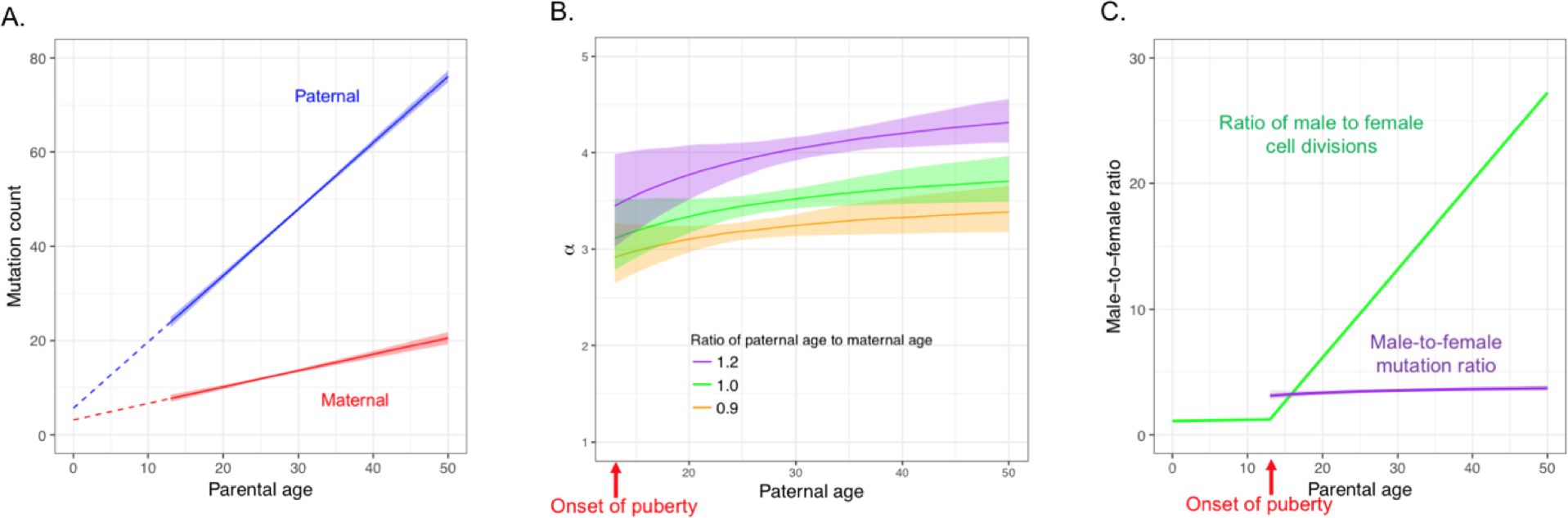
Inferred sex and age dependencies of germline mutations (based on a linear model applied to trios with maternal age no greater than 40 years of age). In all panels, shaded areas and bars represent 95% confidence intervals of the corresponding quantities obtained from bootstrapping. (A) Inferred sex-specific mutation rates as a function of parental ages. Parental ages are measured since birth, i.e., birth corresponds to age 0 (throughout the manuscript). The extrapolated intercepts at age 0 are small but significantly positive for both sexes, implying a weak but significant effect of reproductive age on yearly mutation rates (Gao *et al.*, 2016). (B) Predicted male-to-female mutation ratio (α) as a function of the ratio of paternal to maternal ages. For reference, the ratio of parental ages is centered around 1.10 in the deCODE DNM dataset (s.d.=0.20). (C) Contrast between male-to-female mutation ratio (purple) and the ratio of male-to-female cell divisions (green), assuming the same paternal and maternal ages. Estimates of the cell division numbers for the two sexes in humans are from Drost and Lee (1995).

In addition to a linear model, we considered exponential age effects of either or both sexes. We observed a significant improvement in fit of an exponential maternal age effect over a linear one (ΔAIC = −29.9), consistent with a previous analysis of the Inova dataset that indicated a more rapid increase in the maternal mutation rate at older ages (Wong *et al.*, 2016). To verify our finding, we divided the 1548 trios into two groups with maternal age at conception over or under the median age of 27 years and fit a linear model to the two groups separately. As expected from an accelerating increase in the number of maternal mutations with age, the estimated maternal age effect is greater for older mothers than for younger mothers (0.56 vs 0.24, 95% CI: [0.45,0.66] vs [0.12,0.38]; Table S4), whereas the estimates of paternal age effect are similar for the two groups (1.41 vs 1.40, 95% CI: [1.31, 1.51] vs [1.29, 1.53]). We further found that the exponential maternal age effect no longer provides a significantly better fit when excluding the 72 trios with maternal age over 40 (Table S5). As a sanity check on our estimates, we predicted the paternal mutation fraction for individuals with divergent paternal and maternal ages (G_P_/G_M_=0.9, 1.2 or 1.4); our predictions provide a good fit to the observed patterns for the subset of phased mutations (Fig. S4).

Focusing on the linear model fitted to trios with maternal ages below 40 years at conception (Table S5), the male-to-female mutation ratio is already ∼3 (95% CI: [2.8, 3.5]) at the average age of onset of puberty (assumed to be 13 years of age for both sexes; Fig. 2B,C)(Nielsen *et al.*, 1986), consistent with the observation of stable fraction of paternal mutations with paternal age (Fig. 1), and indicating that the male germline has accumulated a substantially greater number of DNMs than the female germline by puberty. The same is seen in our reanalysis of the smaller Inova dataset (Fig. S5). At face value, this finding is puzzling: male and female germ cells are thought to experience comparable numbers of divisions by then (an estimated 35 vs 31, respectively) and approximately half of these divisions predate sex determination (SD) (Vogel and Rathenberg, 1975; Drost and Lee, 1995), so we would expect males and females to harbor similar numbers of replicative mutations before puberty. Moreover, differences in the mutation spectrum between males and females are subtle (Rahbari *et al.*, 2016; Jónsson, Sulem, Kehr, *et al.*, 2017; I. Agarwal and M. Przeworski, unpublished results), suggesting that the sources of most mutations are shared between the two sexes.

How then to explain that α is already high by puberty and persists at roughly the same value throughout adulthood? Three possible resolutions are that (1) the number of male germ cell divisions from sex determination to puberty has been vastly underestimated; (2) after sex determination, germ cell divisions are much more mutagenic in males than in females; or (3) damage-induced mutations contribute a substantial fraction to male germline mutations by puberty. In principle, the first two possibilities could account for the high α at puberty but they would only lead to a stable α throughout adulthood under implausible conditions. Specifically, the numbers of cell divisions and the mutation rates per cell division over developmental stages in both sexes would need to meet specific conditions, such that male-to-female ratio of replication errors before puberty is coincidentally similar to the ratio of increases in mutation rates in the two sexes after puberty (see Supplementary Materials). Instead, we hypothesize that most germline mutations in both sexes are damage-induced: under this scenario, the elevated α by puberty and the stable α after puberty would reflect damage rates that are roughly constant per unit time in both sexes and higher in males (Ségurel, Wyman and Przeworski, 2014).

### Specific sources of DNA damage-induced mutations

To explore possible mutational mechanisms, following previous studies (Alexandrov *et al.*, 2013; Lindsay *et al.*, 2016; Harris and Pritchard, 2017), we classified point DNMs into six disjoint and complementary mutation classes based on parental and derived alleles: T>A, T>C, T>G, C>A, C>G, and C>T (each type also includes the corresponding variant on the reverse complement strand). Given the well-characterized hypermutability of methylated CpG sites, we further divided the C>T transitions into sub-types in non-CpG and CpG contexts (excluding CpG islands for the latter; see Materials and Methods). Confirming the original analysis of these data (Jónsson, Sulem, Kehr, *et al.*, 2017), we detected significant paternal and maternal age effects for all seven mutation types, of varying magnitudes (Table S1; Fig. S6). While α is stable with parental age for most mutation types (Fig. S6), C>G transversions and C>T transitions at CpG sites stand out from the general pattern (Fig. 3). In particular, C>G mutations show a decreasing α with age and CpG>TpG mutations an increasing α, evident in both the direct analysis of phased mutations and in our modeling of all mutations (Fig. 3A, B). We further found that whereas the linear model is a good fit to the other six mutation types individually, for C>G mutations, a model with a linear paternal age effect and an exponential maternal age effect provides a significantly better fit (ΔAIC = −18.3; Table S3). Interestingly, an exponential maternal age effect also provides a significantly better fit for the six non-C>G point mutations combined (ΔAIC<−9; Table S5), and again the effect disappears when the 72 trios with maternal age over 40 are excluded, suggesting that mutation types other than C>G are also increasing at higher rates in mothers of older ages.

**Figure 3.**
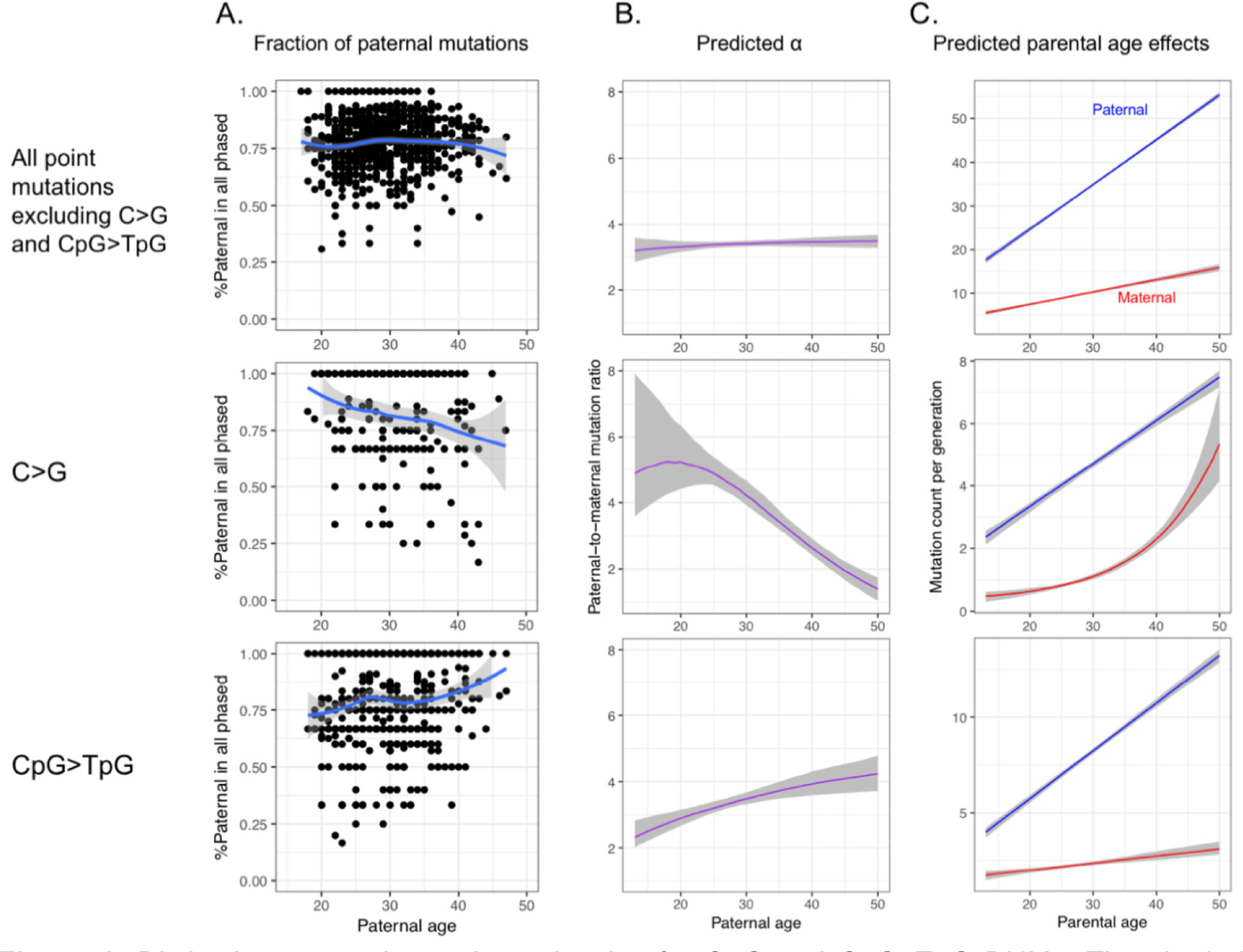
Distinctive sex and age dependencies for C>G and CpG>TpG DNMs. The shaded areas in all panels represent 95% confidence intervals. See Fig. S6 for similar plots for other mutation types. The male-to-female mutation ratio at age 17 is significantly lower for CpG>TpG than for other mutation types (see main text). (A) Fraction of paternal mutations in phased DNMs; (B) Predicted male-to-female mutation ratio (α); (C) Predicted parental age effects.

Based on their spatial distribution in the genome and their increase with maternal age, maternal C>G mutations were hypothesized to be associated with double-strand breaks (DSB) in aging oocytes: this mutation type often appears in clusters with strong strand-concordance and near *de novo* copy number variant breakpoints and is enriched in genomic regions with elevated rates of non-crossover gene conversion-an enrichment that increases rapidly with maternal age (Jónsson, Sulem, Kehr, *et al.*, 2017; Goldmann *et al.*, 2018). C>G mutations are also more frequent in the human pseudo-autosomal 1 region, which experiences an obligate crossover in males, than on autosomes and the rest of the X chromosome (I. Agarwal and M. Przeworski, unpublished results). We additionally found that C>G transversions are significantly more likely than other mutation types to occur on the same chromosome as a *de novo* deletion (≥5bp) in the same individual, and conditional on co-occurrence, that the distance to the closest deletion tends to be shorter (Fig. S7). The same association is not seen for short deletions or insertions (<5bp), however (Table S4), which are more likely to arise from replication slippage (Montgomery *et al.*, 2013; Kloosterman *et al.*, 2015). Together, these observations support imperfect repair of double-strand breaks as an important source of C>G transversions in both sexes (Jónsson, Sulem, Kehr, *et al.*, 2017; Goldmann *et al.*, 2018; I. Agarwal and M. Przeworski, unpublished results).

In turn, the high rates of C>T transitions at CpG sites are thought to be due to the spontaneous deamination of methylcytosine (Lindahl and Nyberg, 1974; Fryxell and Zuckerkandl, 2000) that remains unrepaired by the next replication cycle, though recent studies of tumors indicate that they may also result from an interaction between methylation and the DNA replication process (Poulos, Olivier and Wong, 2017; Tomkova *et al.*, 2018). Since the methylation dynamics of mammalian male and female germlines are relatively well characterized, we can make *a priori* predictions about when sex differences in C>T transitions should arise in development. Specifically, during embryogenesis, several rounds of global DNA demethylation and remethylation take place to enable the erasure and re-establishment of the epigenetic memory from the parents (Reik, Dean and Walter, 2001; Kobayashi *et al.*, 2013). Because these methylation changes are shared by male and female embryos until sex determination of the embryo, the two sexes should share most methylation-related mutations during early development. Consistent with this prediction, we estimated a lower α for CpG transitions than for other presumably more replication-dependent mutation types at early reproductive ages (e.g., at age 17, α=2.6 [2.2, 3.0] vs 3.4 [3.2, 3.7] for mutations other than CpG>TpG) (Fig. 3B). After sex determination (around 7 weeks post fertilization in humans), the methylation profiles of male and female germ cells diverge: re-methylation takes place early in males, before differentiation of spermatogonia, but very late in females, shortly before ovulation (Reik, Dean and Walter, 2001; Kobayashi *et al.*, 2013). Therefore, the male germline is markedly more methylated compared to the female germline for the long period from sex determination of the parents to shortly before conception of the child. Accordingly, after puberty, the estimated yearly increase in CpG>TpG mutations is 6.5-fold higher in fathers than in mothers, roughly double what is seen for other mutation types, resulting in a marked increase in α with parental age at CpG>TpG (Fig. 3C). In support of the key role of methylation in CpG transition mutagenesis, the genomic distribution of DNMs in the 1548 Icelandic trios is strongly associated with the methylation levels in testis of an unrelated male donor in the Roadmap Epigenomics project, and less so with those in ovary of an unrelated female (Fig. S8A)(Lister *et al.*, 2009). Moreover, the distributions of paternal and maternal mutations correspond better to the methylation profiles of testis and ovary, respectively (Fig. S8BC). In summary, the sex and age dependencies of CpG transitions accord with the temporal and genomic methylation profiles of the mammalian germ cells in a sex-specific manner, supporting the notion that methylation-related mutagenic processes are the major sources of CpG transitions, and validating our inferences for the one case in which we have independent information about what to expect. Together with C>G mutations, CpG transitions represent approximately 1/3 of germline mutations accumulated in a parent of age 30 at conception; both show signatures of DNA damage-induced mutational mechanisms.

### The number of *de novo* mutations increases substantially with maternal age

In mammals, primary oocytes are formed and arrested in prophase of meiosis I, before the birth of the future mother, with no further DNA replication occurring until fertilization. On this basis, the maternal age effect detected by recent DNM studies (Goldmann *et al.*, 2016; Wong *et al.*, 2016; Jónsson, Sulem, Kehr, *et al.*, 2017) and confirmed here (Fig. 2A) has been interpreted as reflecting the accumulation of DNA lesions or damage-induced mutations in (primary) oocytes during the lengthy meiotic arrest phase (Ségurel, Wyman and Przeworski, 2014; Gao *et al.*, 2016; Goriely, 2016; Jónsson, Sulem, Kehr, *et al.*, 2017), exemplified by the rapid increase of maternal C>G mutations. However, other explanations for a maternal age effect are possible (Wong *et al.*, 2016). For example, such an effect could also arise if oocytes ovulated later in life have undergone more mitoses (Polani and Crolla, 1991; Fulton *et al.*, 2005). In this scenario, the substantial increase in maternal DNMs from age 17 to age 40 (Fig. 2A) would require oocytes ovulated later in life to go through almost double the number of cell divisions compared to oocytes ovulated early (more, if the per cell division mutation rate is higher in early cell divisions; see below). Moreover, this scenario does not provide an explanation for the stability of the male-to-female ratio with parental ages. Thus, while this phenomenon could hypothetically contribute to the maternal age effect, in practice, it is likely to be a minor effect (see SM for a more detailed discussion).

A more plausible explanation is a maternal age effect on post-zygotic mutations. Although DNMs are usually interpreted as mutations that occur in germ cells of the parents, in fact what are identified as DNMs in trio studies are the genomic differences between the offspring and the parents in the somatic tissues sampled (here, blood). These differences can arise in the parents and also during early development of the child (Fig. 4A; adapted from Moorjani, Gao and Przeworski, 2016). Notably, the first few cell divisions of embryogenesis have been found to be relatively mutagenic, leading to somatic and germline mosaic mutations in a study of four generations of cattle (Harland *et al.*, 2017) and (to a lesser extent) in humans (Huang *et al.*, 2014; Acuna-Hidalgo *et al.*, 2015; Rahbari *et al.*, 2016; Ju *et al.*, 2017; Lim *et al.*, 2017; Sasani *et al.*, 2018), as well as to mutations that are discordant between monozygotic twins (Dal *et al.*, 2014; Jónsson, Sulem, Kehr, *et al.*, 2017). Increased numbers of point mutations in the first few cell divisions should perhaps be expected, as two key components of base excision repair are missing in spermatozoa, leading lesions accumulated in the last steps of spermatogenesis to be repaired only in the zygote (Smith *et al.*, 2013). More generally, mammalian zygotes are almost entirely reliant on the protein and transcript reservoirs of the oocyte until the 4-cell stage (Braude, Bolton and Moore, 1988; Dobson *et al.*, 2004; Zhang *et al.*, 2009). Thus, if the replication or repair machinery of the oocytes deteriorate with the mother’s age (Titus *et al.*, 2013; Wei *et al.*, 2015), one consequence could be more mutations in the first few cell divisions of the embryo (Fig. 4B). This scenario predicts that the mother’s age influences not only the number of mutations on the chromosomes inherited from the mother but also from the father (which would be assigned to “paternal mutations”) (Fig. 4B).

**Figure 4.**
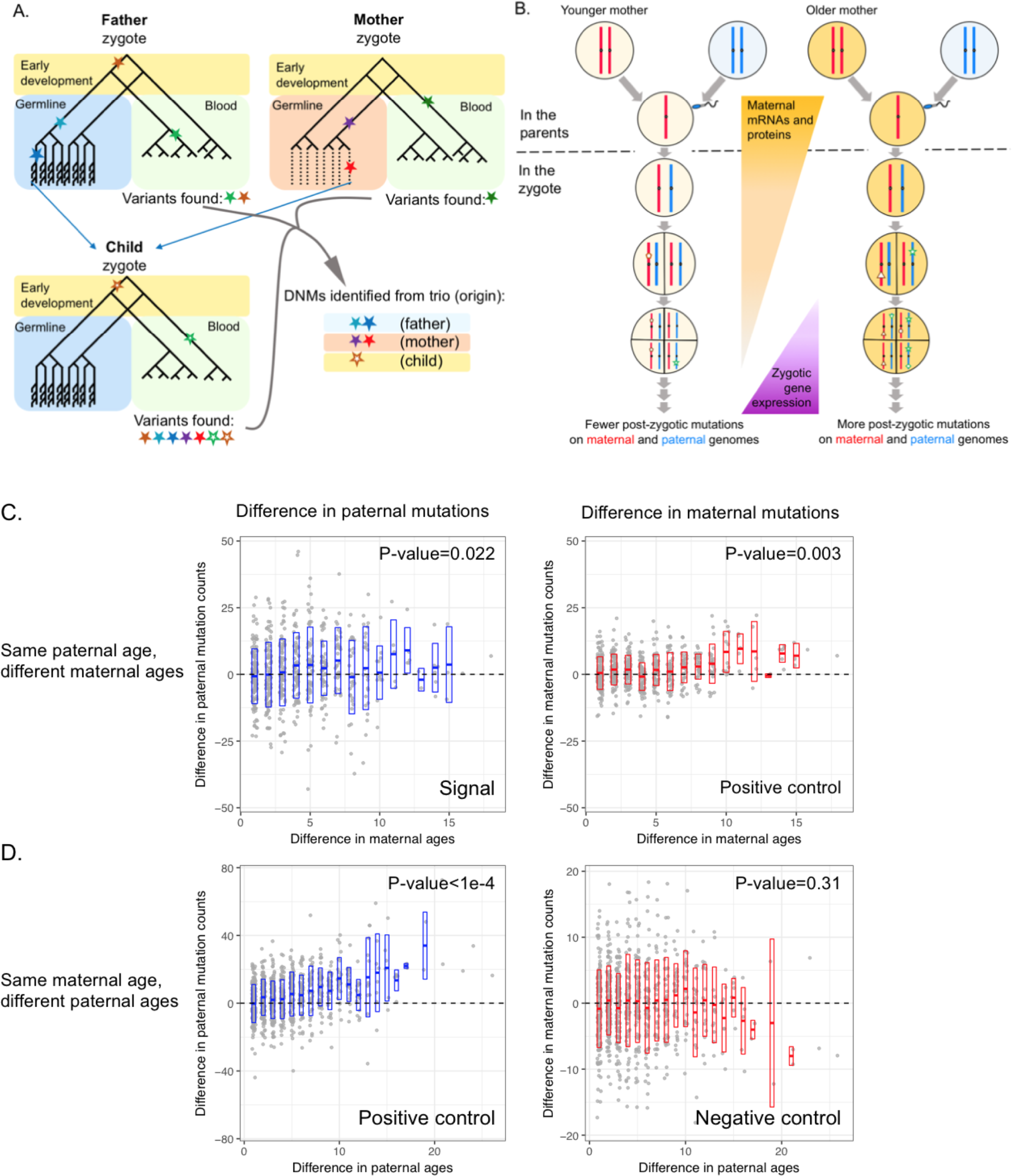
Maternal age effect on mutations that occur on paternally inherited chromosomes. (A) An illustration of mutations occurring during development and gametogenesis (figure adapted from Moorjani, Gao and Przeworski, 2016). In this cartoon, we assume that the most recent common ancestor of all cells in an individual is the fertilized egg. Filled stars represent mutations that arise in the parents and hollow stars in the child. The standard trio approach requires allelic balance in the child and no or few reads carrying the alternative allele in the parent, leading to inclusion of some early post-zygotic mutations in the child (brown hollow) and exclusion of a fraction of early mutations in the parents (brown filled). The two effects partially cancel out when estimating the per generation mutation rate, but potentially lead to underestimate of the fraction of mutations that are early embryonic (Harland *et al.*, 2017). Damage-induced mutations in the oocyte (red filled) and post-zygotic mutations (brown hollow) in the child are both child-specific, but their mosaic status can be evaluated by examining transmissions to a third generation. (B) An illustration of a maternal age effect on the number of post-zygotic mutations. (C) Pairwise comparison conditional on the same paternal age. Each point represents a pair of trios, with x-axis showing the difference in maternal ages and y-axis the difference in paternal mutation counts (left; older mother - younger mother) or maternal mutation counts (right; older mother - younger mother); position is slightly shifted to show overlapping points. P-values are evaluated by 10,000 permutations, using Kendall’s rank correlation test statistic (see Materials and Methods). (D) Pairwise comparison conditional on the same maternal age, similar to (C). The ranges of y-axis differ for the plots on left and right.

Any maternal age effect on post-zygotic mutations is challenging to detect, given the small fraction of post-zygotic mutations estimated in humans(Jónsson, Sulem, Arnadottir, *et al.*, 2017), especially in comparison with the stochasticity in mutation counts across individuals and the noise induced by incomplete phasing of mutations. Further reducing the ability to detect either a pre- or post-zygotic effect of maternal age is the high correlation between maternal and paternal ages (which is why a maternal age effect was not detected in earlier, smaller studies) (Kong *et al.*, 2013; Francioli *et al.*, 2015). As an illustration, if we assume a relatively large increase of 0.3 *paternal* mutations with each year of *maternal* age and complete phasing of DNMs, we estimate the power to detect this maternal age effect in 200 trios to be only 45-56% (Fig. S9). Moreover, using trio data alone, DNMs can only be phased based on informative heterozygous variants in the same reads, so only a small fraction (typically 25-30%) are phased (Goldmann *et al.*, 2016; Wong *et al.*, 2016; Jónsson, Sulem, Kehr, *et al.*, 2017; see Fig. S1). If we assume no DNM calling errors and a uniform phasing rate of 30% across all trios, the power is reduced to 15-20% with 200 trios and is 66-73% even with 1,000 trios (Fig. S9). Moreover, the phasing rate is likely to vary somewhat across families, introducing additional variation in the number of phased mutations and greatly reducing the power. Finally, these simulations ignore errors in calling DNMs, when in reality error rates are non-zero-especially when a third generation is not available to verify transmission-so the power estimates for the standard trio design are likely too high. Consistent with these considerations, we found that in the deCODE dataset, DNMs from trios with and without third-generation individuals differ in multiple properties, including the dependency of the number of mutations on sex and parental age (see SM). Therefore, we expect that a maternal age effect on post-zygotic mutations, if it exists, should only be detectable using data from a sufficiently large number of multi-generation pedigrees. To our knowledge, the only publicly available dataset that currently satisfies these criteria is the deCODE dataset, which includes 225 three-generation families(Jónsson, Sulem, Kehr, *et al.*, 2017).

To reduce noise due to incomplete phasing and DNM calling errors, we focused on the 199 deCODE probands in which almost all DNMs are phased (>95% phased) by transmission to third-generation individuals. Intriguingly, Poisson regression of the count of paternal mutations on both parental ages revealed an effect of maternal age (p=0.035) and a slight but nonsignificant improvement in the fit compared to a model with paternal age only (ΔAIC=−2.4; Table S7). We verified that such a signal would not arise artifactually from the correlation between maternal and paternal ages and the assignment of parental ages to 1-year bins (p=0.007; see Materials and Methods for details). We further noted that the maternal age effect becomes more significant when we limited our analysis to 130 probands with >98% DNMs phased (p=0.004). To visualize the effect, we carried out further analyses of DNMs in the 199 probands by comparing all pairs with the same paternal age but different maternal ages. Among 619 such pairs, the child born to the older mother carries more paternal mutations than does the child with the younger mother in 319 cases, fewer in 280 cases, and the same number in 20 cases, and greater differences in the number of paternal mutations are associated with greater differences in maternal ages (Kendall’s rank test τ=0.09, *p*=0.024 by a permutation test; see Materials and Methods for details; Fig. 4C). In contrast, there is no significant effect of paternal age on the number of maternal mutations when matching for the mother’s age (p>0.31; Fig. 4D; Table S7), although the power to detect such an effect is limited because of the smaller numbers of maternal mutations.

The estimated effect of maternal age on maternal mutations is 0.34 mutations per year (s.e.=0.04) by Poisson regression (p=3.4e-13). The estimated maternal age effect on paternal mutations is similar but highly uncertain (0.30, s.e.=0.14). Naively, one might expect the maternal age effect on maternal mutations to be stronger, as it includes both pre-zygotic effects (e.g., damage in the oocyte) and post-zygotic effects, whereas the effect on paternal mutations can only be post-zygotic. This expectation is implicitly based on the assumption of same post-zygotic effects of maternal age on maternal and paternal genomes, but they need not be. Indeed, before fertilization, sperm and oocytes may harbor different levels of DNA damage (e.g., oxidative stress may be higher in male germ cells) (De luliis *et al.*, 2009; Lim and Luderer, 2011) and after fertilization but before the first cleavage, the two parental genomes experience distinct epigenetic remodeling and are replicated separately, in their own pronuclei (Ferreira and Carmo-Fonseca, 1997; Mayer *et al.*, 2000). Thus, even if the precise effects of maternal age on the zygote were known, the relative contributions of pre-zygotic and post-zygotic effects of maternal age on the maternal genome are not distinguishable without additional data. Regardless, the positive association between maternal age and the number of DNMs on paternal chromosomes supports the hypothesis that a mother’s age at conception affects the post-zygotic mutation rate in the developing embryo.

In cattle and humans, the high frequency mosaic mutations that are likely to have arisen in early embryonic development are enriched for C>A transversions (Huang *et al.*, 2014; Harland *et al.*, 2017; Ju *et al.*, 2017), potentially reflecting the accumulation of the oxidative DNA damage 8- hydroxyguanine in oocytes and the last stages of spermatogenesis (De luliis *et al.*, 2009; Lim and Luderer, 2011) that remains uncorrected in spermatozoa (Smith *et al.*, 2013). Hypothesizing that a maternal age effect may be particularly pronounced for these mutations, we focused on C>A mutations in the 199 probands with high phasing rates. Although this subset represents only ∼8% of mutations, there is a significant effect of maternal age on paternal mutations (p=0.02 by Poisson regression), and the point estimate of the maternal age effect on paternal genome (0.095, s.e.=0.041) is even stronger than that of the paternal age (0.057, s.e.=0.033), as well as stronger than the maternal age effect on maternal genome (0.024, s.e.=0.0094; see Materials and Methods for details). Such results are not often obtained in a random subset of mutations of the same size (p=0.045; see Materials and Methods), suggesting paternal C>A mutations are indeed more strongly affected by maternal age than are other DNMs. After excluding C>A mutations, the maternal age effect on paternal point mutations is no longer significant in the 199 probands with >95% DNMs phased (p=0.13) but remains significant for the subset of 130 probands with phasing rates higher than 98% (p=0.02). Thus, a mutation type associated with damage in sperm and known to be enriched in early embryogenesis shows a heightened signal of a maternal age effect on paternal mutations, without entirely accounting for the signal.

## Discussion

These findings call into question the textbook view that germline mutations arise predominantly from replication errors in germ cells. First, multiple lines of evidence suggest that CpG transitions and C>G mutation often arise from methylation-associated damage and doublestrand break repair, respectively. Second, even excluding both of these mutation types, roughly three-fold more paternal mutations than maternal mutations have occurred in very young parents, despite similar numbers of estimated germ cell divisions by that age (Fig. 2B, C). Moreover, the male-to-female mutational ratio remains surprisingly stable with parental age, even as the ratio of male-to-female cell divisions increases rapidly (Fig. 2B, C). The high α of ∼3 in young parents could be explained by a vast underestimation of the number of germ cell divisions in males between birth and puberty, but its stability with parental ages cannot. Lastly, despite highly variable cell division rates over development, germline mutations accumulate in rough proportion to absolute time in each sex (Fig. 2A; Fig. S6). Together, these findings point to a substantial role of DNA damage-induced mutations, raising questions about the relative importance of endogenous versus exogenous mutagens, as well as about why male and female germ cells differ in the balance of DNA damage and repair.

We further report on tentative signals of a maternal age effect on paternal mutations, which support the hypothesis that the age of a mother at conception influences mutagenesis in early embryonic development of her child. Because violations of the assumptions of Poisson regression may lead to inflated significance and the various signals that we detected were based on analyses of the same data, our current findings need to be replicated in large, independent datasets. The hypothesis of a maternal age effect on the post-zygotic mutation rate appears plausible, however, given what is known about early mammalian embryogenesis and accumulating evidence for a non-negligible number of early embryonic mutations among DNMs (Lindsay *et al.*, 2016; Rahbari *et al.*, 2016; Harland *et al.*, 2017; Jónsson, Sulem, Arnadottir, *et al.*, 2017; Sasani *et al.*, 2018), and thus will be important to investigate further.

Our findings also carry implications for the differences in mutation rates that have been reported across populations and species (e.g., Hwang and Green, 2004; Moorjani *et al.*, 2016). Within mammals, an older age of reproduction of a species is associated with a decreased ratio of X to autosome divergence (Makova and Li, 2002; Wilson-Sayres *et al.*, 2011)(interpreted as higher male-to-female mutation ratios; but see Amster and Sella, 2016; an older age of reproduction is also correlated with a lower substitutions rate, with a much weaker relationship seen for CpG transitions (Li and Tanimura, 1987; Kim *et al.*, 2006; Ségurel, Wyman and Przeworski, 2014; Moorjani *et al.*, 2016). These observations have been widely interpreted as supporting a replicative origin of most non-CpG transitions (Li *et al.*, 1996; Kim *et al.*, 2006; Wilson-Sayres *et al.*, 2011; Ségurel, Wyman and Przeworski, 2014; Moorjani *et al.*, 2016). Instead, our analyses of human DNMs show generally weak effects of reproductive age on the male-to-female mutation ratio as well as on yearly mutation rates (Fig. S6BC), an important role for non-replicative mutations beyond CpG transitions, and a potential coupling of maternal and paternal age effects. One possibility is that inter-species differences in the male-to-female mutation ratio and in substitution rates instead reflect changes in the ratio of paternal and maternal ages at reproduction (Fig. 2B)(Amster and Sella, 2016) and in rates of DNA damage (e.g., metabolic rates) that covary with life history traits (Martin and Palumbi, 1993; Wilson-Sayres *et al.*, 2011).

## Materials and Methods

### Processing of *de novo* mutation data

For each *de novo* mutation (DNM), we obtained parental ages at conception of the child (proband) and the position, allele and parent-of-origin information from the Supplementary Material of the publication for one dataset (Jónsson, Sulem, Kehr, *et al.*, 2017) and by personal communication with the authors for the replication dataset (Goldmann *et al.*, 2016; Wong *et al.*, 2016). We considered a mutation as “phased” if the parental haplotype on which it arose was determined by either informative flanking variant in the read or from transmission to a third generation. See Table S2 for a comparison of summary statistics of these datasets.

For both datasets, we removed indels and mutations on X chromosome (no Y-linked DNMs were reported), which resulted in 98,858 and 35,793 point mutations (or single nucleotide substitutions) for Jónsson, Sulem, Kehr, *et al.* (2017) and Goldmann *et al.* (2016), respectively. Each of these mutations was then assigned into one of six mutation types (T>A, T>C, T>G, C>A, C>G, and C>T) based on the original allele present in homozygous state in both parents and the derived allele that is carried by the child in heterozygous state. Complementary combinations (such as C>T and G>A) were combined such that the original allele is always a pyrimidine (C or T). Moreover, each DNM was annotated to be in CpG or nonCpG context based on its two immediate flanking bases extracted from human reference genome. For analyses of C>T mutations at CpG sites, we excluded CpG transitions present in CpG islands (annotations downloaded from UCSC browser: CpG Islands track), because these sites are thought to be hypomethylated and thus behave differently in terms of mutation rate compared to CpG sites outside CpG islands (CGI) (Moorjani et al. 2016). C>T mutations at CpG sites in CGIs were included in analysis of “all point mutations”.

### Estimation of sex-specific mutation parameters with a model-based approach

Similar to Jónsson, Sulem, Kehr, *et al.* (2017), we modeled the expected number of mutations from a parent as a linear function of her (or his) age at conception of the child, and assumed that the observed maternal (paternal) mutation count follows a Poisson distribution with this expectation. One difference from the Jónsson, Sulem, Kehr, *et al.* (2017) model is in how we account for the incomplete parental origin information for the unphased DNMs. Unlike Jónsson et al., we explicitly modeled the phasing process as a binomial sampling of DNMs with a proband-specific phasing rate parameter, assuming that the phasing probabilities of all mutations in the same individual are identical and independent (as seems sensible). This approach enabled us to fully leverage information from both phased and unphased mutations jointly.

Specifically, the increase in DNMs with sex-specific parental ages is modeled as the following:

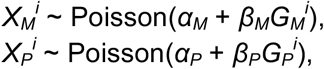

where index *i* indicates the proband; 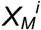 and 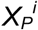 are the total numbers of maternal and paternal mutations; 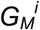 and 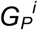 are ages of the mother and the father at conception, respectively; *α*_*M*_, *β*_*M*_, *α*_*P*_, and *β*_*P*_ are the mutation parameters that characterize the sex-specific parental age effects and are shared across all probands (note that *α*_*M*_ and *α*_*P*_ are the extrapolated intercepts at age zero, which are not necessarily non-negative). We assumed linear effects for both sexes in the initial model, but relaxed this assumption by testing for exponential effects for either or both sexes later (see “Test for alternative models for parental age effects” below; Table S3).

Because of incomplete phasing, 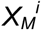 and 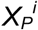 are not directly observed. Thus, we modeled the observed mutation counts as:

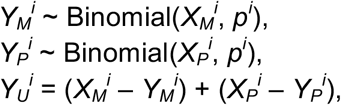

where *p*^*i*^ is the phasing rate in proband *i* and 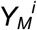, 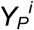 and 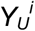 represent the numbers of phased maternal, phased paternal and unphased mutations, respectively. 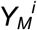, 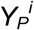 and 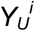 are defined as random variables, and we denote the observed values of these with lower case notations 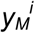, 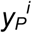 and 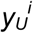.

With the parameterization above, the likelihood of the observed data for proband *i* can be written as:

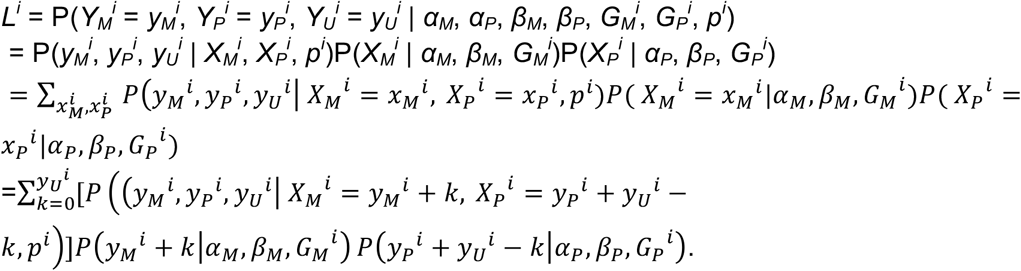

We note that the likelihood function of Jónsson et al. (2017) does not include the first term, which is the probability of the observed data given possible partitionings of the unphased mutations into paternal and maternal origins (assuming the same phasing rates of maternal and paternal mutations). As an illustration, the set of observations 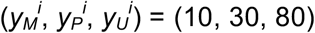 is more probable under 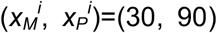, where one third of DNMs were phased for both parental origins, than under 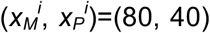, where 75% paternal DNMs were phased but only 12.5% of maternal DNMs.

The likelihood function for proband *i* can be simplified as (see “Derivation of the likelihood function for estimating sex-specific mutation parameters” section in SM):

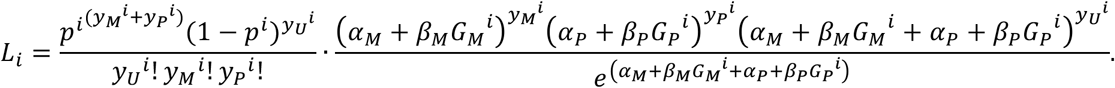

Note that the first term contains the phasing rate (*p*^*i*^) but is independent of the mutation parameters, whereas the second term is dependent on the mutation parameters but independent of *p*^*i*^ Therefore, the maximum likelihood estimator (MLE) of *p*^*i*^ and those of the mutation parameters can be identified by maximizing the first and second terms separately.

The log joint likelihood of all observed data under a set of mutation parameter values can be expressed as:

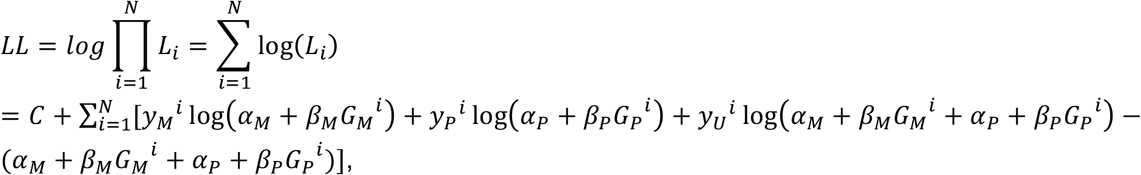

where *C* is a constant that is independent of the mutation parameters of interest.

We implemented this log likelihood function in R and found the MLEs of the mutation parameters by using function mle2 in the package “bbmle” (with the optimization method “BFGS”; Bolker, R Development Core Team 2017). To avoid being trapped in local maxima, we tested a grid of initial values for the slopes (*β*_*P*_ and *β*_*M*_). We performed the estimation for all point mutations altogether as well as for each mutation type separately.

### Confidence intervals of male-to-female mutation ratio at given parental ages

To account for uncertainties in the DNM parameter estimates, we used a bootstrap approach, randomly re-sampling the probands with replacement 500 times, keeping the same total number of probands in each run. For each replicate, we obtained the MLEs of the DNM parameters as described above, predicted the numbers of paternal and maternal mutations at given ages, and calculated the male-to-female mutation ratio; the actual average ages at conception in the Icelandic dataset are 28.2 and 32.0 for mothers and fathers, respectively). Thus, each bootstrap provides one point estimate for each of the quantities of interest, and the approximate distribution for each quantity can be obtained by aggregating results from the 100 replicates.

### Test for alternative models for parental age effects

In addition to the linear model described in the above, we also considered models with exponential parental age effects post-puberty for either or both sexes. Specifically, we modeled the exponential parental age effect as follows:

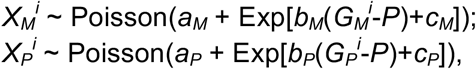

where *P*=13 is the age of onset of puberty assumed for both sexes. We note that results are not sensitive to the choice of the value of *P*. Under this formulation, models with different values of *P* are mathematically equivalent to models with the same *b*_*P*_ (or *b*_*M*_) value but different *c*_*P*_ (or *c*_*M*_) values. Indeed, we confirmed the MLEs for *b*_*P*_ and *b*_*M*_ are the same for different *P* values (even for *P*=0).

We obtained the MLEs and corresponding log likelihoods of all four models for all point mutations combined and for each mutation type separately, and used the Akaike information criterion (AIC) to compare the relative fits of different models (a smaller AIC indicates a better fit of the model). We took ΔAIC<−6 as the threshold for evidence for a significantly better fit (approximately 20-fold more probable). The models with exponential paternal age effect provide worse fits (ΔAIC>0) for all mutation types.

For all DNMs combined, models with exponential effects of maternal age or both parental ages provide significantly better fits but are not significantly different from each other (Table S3). As verification, we split the 1548 trios into two groups with maternal age at conception over and under 27 years (the median maternal ages in the dataset), respectively, and fitted both with linear parental age effects. The estimate of the maternal age effect is greater for older mothers than for younger mothers (0.56 vs 0.24, 95% CI: [0.45,0.66] vs [0.12,0.38]), whereas the estimates of paternal age effect are similar for the two groups (1.41 vs 1.40, 95% CI: [1.31, 1.51] vs [1.29, 1.53]; Table S3). The improved fit by an exponential maternal age was no longer significant when excluding the 72 trios with maternal age above 40 (Table S5). We therefore considered the linear model fitted to trios with maternal ages under 40 years for analyses for all point mutations combined throughout the manuscript. The predictions based on linear models fitted to all trios and trios with Gm under 40 show similar results: for instance, the estimated male-to-female mutation ratio at puberty (*P*=13) is 3.1 vs 3.3 (95% CI [2.8, 3.5] vs [3.0, 3.7]).

Among all mutation types considered, C>G transversions are the only type for which the model with exponential maternal age effect provides a significantly better fit by the criterion of ΔAIC<−6 (Table S3). Therefore, in all analyses for C>G transversions (e.g., calculation of α), we used the estimates from the model with an exponential maternal age effect (and linear paternal age effect) fitted to all 1548 trios. For all DNMs combined, the model with an exponential maternal age effect also provides a significantly better fit than the linear model. Interestingly, considering all trios, even after C>G transversions (or C>G transversions and CpG transitions) are excluded, an exponential maternal age effect still provides a significantly better fit for other point mutations combined (ΔAIC<−9; Table S5), suggesting that the signal is not driven by C>G mutations alone. Again, this effect is no longer discernible when trios with maternal age above 40 are excluded (Table S4).

### Processing of ovary and testis methylation data at CpG sites

The methylation data were generated as part of the Roadmap Epigenomics project (Lister *et al.*, 2009). We downloaded the methylation data from GEO (https://www.ncbi.nlm.nih.gov/geo/) with accession numbers GSM1010980 for ovary, and GSM1127119 for testis (sperm). The methylation levels were measured by bisulfate sequencing of testis spermatozoa primary cells from a male donor (age and descent unknown) and ovary cells from a 30-year-old female donor of European descent, respectively. Methylation levels (measured as percentage methylated) are reported only for CpG sites in the reference human genome: 27,057,581 (94.3%) of the ∼28,700,000 CpG sites (∼57,400,000 basepairs) have reported methylation levels in ovary, and 26,693,016 (93%) have that information for testis. CpG sites with data available were sorted based on their modification levels and grouped into bins of 100,000 sites (i.e., 50,000 CpGs). The average methylation level of each bin was then correlated with the total number of C>T DNMs in the 1548 Icelandic trios that occurred at the 100,000 sites (an estimate of the average mutation rate of these sites). We note that the methylation profile of ovary cells may be a poor proxy for that of (primary) oocytes, so the correlation between CpG>TpG DNM rate and methylation may be under-estimated. In addition, there is likely inter-individual variation in methylation profiles, but such variation is typically smaller than inter-tissue variation (Schultz *et al.*, 2015), so it is expected to reduce the correlation between methylation and mutation rates in both tissues by a small amount.

### Estimate of the power to detect a maternal age effect on the paternal mutation rate by simulation

We simulated paternal mutation counts for various paternal (y-axis) and maternal (x-axis) age effect sizes on paternal mutation rate, assuming that the mutation count is Poisson distributed, and quantified the fraction of simulations with a significant maternal age effect in 1,000 replicates. We assumed that the extrapolated intercept at *G*_*P*_=0 and *G*_*M*_=0 is six mutations in the assayable regions of a diploid genome (estimated to be 5.56 by deCODE and 6.05 by our analysis), i.e., 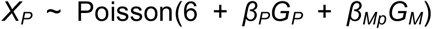. We performed simulations under different combinations of parental effect sizes: 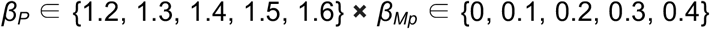 with a sample size of 100, 200, 500, 1000 or 2000 trios. The criteria for a significant maternal age effect are (1) that the fit of a model with both parental ages is improved compared to a model with paternal age only by at least Δ AIC=−2.4; and (2) that the estimated effect of maternal age is positive and the p-value is smaller than 0.05 in the model with both parental ages. We assumed no error in DNM calling, when error rates are in fact non-zero and post-zygotic mutations in particular are more likely to be missed. For the case of incomplete phasing, we took a phasing rate of 0.3, which is the value typically obtained for whole-genome trio data based on informative heterozygous sites in reads, and assumed identical and independent phasing probabilities across mutations and trios, regardless of parental origin. Based on the thinning property of Poisson distribution, the number of phased paternal mutations also follows a Poisson distribution with the product of the mutation rate and the phasing rate as the rate parameter. However, the actual phasing rate is likely to be variable across trios (depending on trio-specific sequencing coverage, and other factors), which will introduce additional variation in the counts of phased mutations and further reduce the power. Given the overly optimistic assumptions about DNM calling and phasing in the simulations, the true power to detect a maternal age effect on paternal mutations is likely lower. As expected, the power to detect a maternal age effect on paternal mutations increases with the simulated effect size, phasing rate and the sample size. See main texts for a brief description of the results.

### Detection and estimation of maternal age effect on paternal mutation rate

For analyses in this section, we focused on the 199 probands in which almost all DNMs were phased (>95% DNM phased). We first did a Poisson regression (with an identity link) of the count of paternal point mutations on both parental ages and found a significant effect of the maternal age (p= 0.035) and a slight but non-significant improvement in the fit compared to a model with paternal age only (ΔAIC=−2.4; approximately 3.3-fold more probable); p-values and AIC obtained by glm() function in R (with option “family = poisson(link = “identity”)”). In contrast, regressing the maternal mutation count on both parental ages does not provide any improvement in the fit compared to a model with maternal age alone (ΔAIC=0.2), although the power to detect an effect of paternal age on maternal mutation, if any, is lower due to the low mutation counts. See Table S7 for estimated effect sizes by Poisson regression.

Motivated by this finding, we re-estimated the mutation parameters by maximum likelihood under models including a maternal age effect on paternal mutations (i.e., “maternal-on-paternal effect”) of the same size (model 1) or a different size (model 2) than the maternal age effect on maternal mutations. Both models provide slight but insignificant improvements in fit compared to a model without a maternal age effect on paternal mutations (model 0) (Table S7), and the model with the same maternal effect on both maternal and paternal mutations gives the best fit based on AIC (ΔAIC =−3.7; MLE of maternal age effect is 0.34 mutations per year; Table S7).

We then carried out two types of analyses of the same data, conditional on paternal age. First, we performed “pairwise analysis”, in which we compared all pairs of trios with the same paternal age, *G*_*p*_ but different maternal ages, *G*_*m*_ (i.e., a “pairwise analysis”). In 619 such pairs, the child born to the older mother carries more paternal mutations than does the child with the younger mother in 319 cases, fewer in 280 cases, and the same number in 20 cases, and the difference in the number of paternal mutations is significantly associated with the difference in the maternal age (Kendall’s rank test tau=0.09, p= 0.0015). Because some of the pairs include the same probands and are thus not independent, we did a permutation test by swapping the maternal ages within paternal age bin and calculating the adjusted z-score of Kendall’s tau-b statistic. 220 out of 10,000 permutations had statistics equal to or greater than that observed with in real data (corresponding to an empirical one-tailed p-value of 0.022). To estimate the effect size of maternal age, we ran weighted linear regression of the difference in paternal counts on the difference in maternal ages for each pair of trios with the same paternal age (with an intercept of zero), with the weight of each data point specified as the inverse of the paternal age, which is approximately proportional to the variance in the observed difference in paternal mutation counts (Table S7). Because the mutation counts are integers and do not follow a normal distribution, the standard errors are inaccurate.

One concern is that parental ages are assigned to integer bins in the Icelandic dataset, and there is potentially a subtle correlation between maternal and paternal ages even within a paternal age bin, in which case variation in paternal age counts caused by small *G*_*P*_ variation may be mistakenly ascribed to an effect of *G*_*M*_. To address this concern, we simulated data of 199 trios with similar parental age structure but no maternal age effect on paternal DNMs, and asked how frequently analysis of simulated data based on binned parental ages would generate signals comparable to those observed in actual data. To mimic the distributions of maternal and paternal ages and the correlation between them in the actual dataset, we simulated an exact maternal age for each trio by adding a random variable that is uniform on (0,1) to the integer maternal age given in the dataset, and a corresponding exact paternal age taken from 2.70 + 1.076*G*_*M*_ + *e*, where *e* follows Normal(0, 4.5) (parameters obtained by ordinary linear regression on the binned parental ages in the dataset). We then simulated the paternal DNM count as a Poisson random variable with expectation of either 1.51*G*_*P*_+6.05 (as estimated by Jónsson, Sulem, Kehr, *et al.*, 2017) or 1.41*G*_*P*_+5.56 (estimated by our maximum likelihood model) and ran Poisson regression or pairwise analysis on the mutation counts and integer parts of parental ages, as described above. The simulated data generated either a greater or equal maternal age effect on paternal mutations by Poisson regression or a more z-score of Kendall’s tau-b statistic as significant or more significant in only about 3.5% of 10,000 replicates and both in ∼0.7%, suggesting that this scenario is unlikely to lead to the patterns observed (Table S8).

Although C>A mutations only constitute 8% of all DNMs, for this mutation type, we found a significant effect of the maternal age (p= 0.020) and a slight improvement in the fit compared to a model with paternal age only (ΔAIC=−3.05; approximately 4.6-fold more probable) by Poisson regression (with identity link) of the number of paternal mutations. More surprisingly, the point estimate of the maternal age effect on paternal genome (0.095; se=0.041) is even stronger than that of the paternal age (0.057, se=0.033) and also stronger than the effect of maternal age on maternal genome (0.024, se=0.0094 by Poisson regression of maternal mutations on maternal age). To test the significance of this finding, we used simulations to examine whether the observations of C>A can happen by chance, conditional on the maternal age effect on paternal mutations on overall DNMs. We focused on 199 trios with >95% DNMs phased and simulated data with two schemes (1) randomly subsampling 8.3% paternal DNMs as C>A mutations for each trio, and (2) shuffling the mutation type labels across all paternal DNMs of the 199 trios. We then ran Poisson regression on the simulated paternal C>A mutation counts and found that in only 4.5% of the 20,000 replication, the model with maternal age would provide a better fit with ΔAIC<-3 and a greater point estimate of maternal age effect than paternal age effect (Table S9). These results suggest that paternal C>A mutations are more strongly affected by maternal age compared to other DNMs. In addition, the fraction of C>A DNMs is higher among paternal mutations than maternal ones (constituting 8.3% of paternal DNMs vs 6.2% of maternal ones) (Jónsson, Sulem, Kehr, *et al.*, 2017), potentially reflecting DNA oxidative stress in spermatogenesis and lack of a complete base excision repair pathway in spermatozoa (De Iuliis *et al.*, 2009; Smith *et al.*, 2013). We also noted that this mutation type is 15-20% underrepresented in the deCODE data set relative to what we expect from an analysis rare variants (present in 3-9 copies) in 7,509 non-Finish Europeans in the gnomAD dataset, after accounting for GC content and effect of GC-biased gene conversion (Z. Gao, P. Moorjani and M. Przeworski, unpublished results), potentially reflective of an under-detection of early embryonic mutations by the standard trio approach. For C>A mutations in the 199 probands with >95% DNMs phased, we did not observe enrichment in the GCA or TCT trinucleotide context reported in Harland et al., possibly due to lack of power.

## Acknowledgments

We thank I. Agarwal, M. Eisen, A. Harpak, J. Pritchard, Z. Williams and members of the Przeworski and Pritchard labs for helpful discussions, as well as G. Coop’s lab, Chuck Langley, Wendy Wong and Vladimir Seplyarskiy for comments on an earlier version of the manuscript.

## References

Acuna-Hidalgo, R. et al. (2015) ‘Post-zygotic Point Mutations Are an Underrecognized Source of de Novo Genomic Variation’, American Journal of Human Genetics. The American Society of Human Genetics, 97(1), pp. 67–74. doi: 10.1016/j.ajhg.2015.05.008.

Acuna-Hidalgo, R., Veltman, J. A. and Hoischen, A. (2016) ‘New insights into the generation and role of de novo mutations in health and disease’, Genome Biology. Genome Biology, 17(1), pp. 1–19. doi: 10.1186/s13059-016-1110-1.

Alexandrov, L. B. et al. (2013) ‘Signatures of mutational processes in human cancer’, Nature, 500(7463), pp. 415–421. doi: 10.1038/nature12477.

Amster, G. and Sella, G. (2016) ‘Life history effects on the molecular clock of autosomes and sex chromosomes’, Proceedings of the National Academy of Sciences, 113(6), pp. 1588–1593. doi: 10.1073/pnas.1515798113.

Braude, P., Bolton, V. and Moore, S. (1988) ‘Human gene expression first occurs between the four- and eight-cell stages of preimplantation development’, Nature, 332(6163), pp. 459–61.

Crow, J. F. (1997) ‘The high spontaneous mutation rate: is it a health risk?’, Proceedings of the National Academy of Sciences of the United States of America, 94(16), pp. 8380–6. doi: 10.1073/pnas.94.16.8380.

Crow, J. F. (2000) ‘The origins, patterns and implications of human spontaneous mutation’, Nature Reviews Genetics, 1(1), pp. 40–47. doi: 10.1038/35049558.

Dal, G. M. et al. (2014) ‘Early postzygotic mutations contribute to de novo variation in a healthy monozygotic twin pair’, Journal of Medical Genetics, 51(7), pp. 455–459. doi: 10.1136/jmedgenet-2013-102197.

Dobson, A. T. et al. (2004) ‘The unique transcriptome through day 3 of human preimplantation development’, Human Molecular Genetics, 13(14), pp. 7–9. doi: 10.1093/hmg/ddh157.

Drost, J. B. and Lee, W. R. (1995) ‘Biological Basis of Germline Mutation : Comparisons of Spontaneous Germline Mutation Rates’, Environmental and molecular mutagenesis, 64(1 995), pp. 48–64.

Ferreira, J. and Carmo-Fonseca, M. (1997) ‘Genome replication in early mouse embryos follows a defined temporal and spatial order.’, Journal of cell science, 110 (Pt 7, pp. 889–897.

Francioli, L. C. et al. (2015) ‘Genome-wide patterns and properties of de novo mutations in humans’, Nature Genetics. Nature Publishing Group, 47(7), pp. 822–826. doi: 10.1038/ng.3292.

Fryxell, K. J. and Zuckerkandl, E. (2000) ‘Cytosine deamination plays a primary role in the evolution of mammalian isochores’, Molecular Biology and Evolution, 17(9), pp. 1371–1383. doi: 10.1093/oxfordjournals.molbev.a026420.

Fulton, N. et al. (2005) ‘Germ Cell Proliferation and Apoptosis in the Developing’, 90(May), pp. 4664–4670. doi: 10.1210/jc.2005-0219.

Gao, Z. et al. (2016) ‘Interpreting the Dependence of Mutation Rates on Age and Time’, PLoS Biology, 14(1), pp. 1–16. doi: 10.1371/journal.pbio.1002355.

Goldmann, J. M. et al. (2016) ‘Parent-of-origin-specific signatures of de novo mutations’, Nature Genetics, 48(8), pp. 935–939. doi: 10.1038/ng.3597.

Goldmann, J. M. et al. (2018) ‘Germline de novo mutation clusters arise during oocyte aging in genomic regions with high double-strand-break incidence’, Nature Genetics. Springer US, pp. 1–6. doi: 10.1038/s41588-018-0071-6.

Goriely, A. (2016) ‘Decoding germline de novo point mutations’, Nature Genetics. Nature Publishing Group, 48(8), pp. 823–824. doi: 10.1038/ng.3629.

Harland, C. et al. (2017) ‘Frequency of mosaicism points towards mutation-prone early cleavage cell divisions’, bioRxiv.

Harris, K. and Pritchard, J. K. (2017) ‘Rapid evolution of the human mutation spectrum’, eLife, 6, pp. 1–17. doi: 10.7554/eLife.24284.

Huang, A. Y. et al. (2014) ‘Postzygotic single-nucleotide mosaicisms in whole-genome sequences of clinically unremarkable individuals’, Cell Research. Nature Publishing Group, 24(11), pp. 1311–1327. doi: 10.1038/cr.2014.131.

Huttley, G. A. et al. (2000) ‘How important is DNA replication for mutagenesis?’, Molecular Biology and Evolution, 17(6), pp. 929–937. doi: 10.1093/oxfordjournals.molbev.a026373.

Hwang, D. G. and Green, P. (2004) ‘Bayesian Markov chain Monte Carlo sequence analysis reveals varying neutral substitution patterns in mammalian evolution’, proceedings of the National Academy of Sciences, 101(39), pp. 13994–14001. doi: 10.1073/pnas.0404142101.

De luliis, G. N. et al. (2009) ‘DNA Damage in Human Spermatozoa Is Highly Correlated with the Efficiency of Chromatin Remodeling and the Formation of 8-Hydroxy-2′-Deoxyguanosine, a Marker of Oxidative Stress1’, Biology of Reproduction, 81(3), pp. 517–524. doi: 10.1095/biolreprod.109.076836.

Jónsson, H., Sulem, P., Kehr, B., et al. (2017) ‘Parental influence on human germline de novo mutations in 1,548 trios from Iceland’, Nature. Nature Publishing Group, 549(7673), pp. 519–522. doi: 10.1038/nature24018.

Jónsson, H., Sulem, P., Arnadottir, G., et al. (2017) ‘Recurrence of de novo mutations in families’, bioRxiv.

Ju, Y. S. et al. (2017) ‘Somatic mutations reveal asymmetric cellular dynamics in the early human embryo’, Nature. Nature Publishing Group, 543(7647), pp. 714–718. doi: 10.1038/nature21703.

Kim, S. H. et al. (2006) ‘Heterogeneous genomic molecular clocks in primates’, PLoS Genetics, 2(10), pp. 1527–1534.

Kloosterman, W. P. et al. (2015) ‘Characteristics of de novo structural changes in the human genome’, Genome Res., 25, pp. 792–801. doi: 10.1101/gr.185041.114.19.

Kobayashi, H. et al. (2013) ‘High-resolution DNA methylome analysis of primordial germ cells identifies gender-specific reprogramming in mice’, Genome Research, 23(4), pp. 616–627. doi: 10.1101/gr.148023.112.

Kong, A. et al. (2013) ‘Rate of de novo mutations, father’s age, and disease risk’, Nature, 488(7412), pp. 471–475. doi: 10.1038/nature11396.Rate.

Kumar, S. and Subramanian, S. (2002) ‘Mutation rates in mammalian genomes’, proceedings of the National Academy of Sciences, 99(2), pp. 803–808. doi: 10.1073/pnas.022629899.

Li, W.-H. et al. (1996) ‘Rates of nucleotide substitution in primates and rodents and the generation-time effect hypothesis’, Molecular phylogenetics and Evolution, 5(1), pp. 182–187. doi: 10.1006/mpev.1996.0012.

Li, W.-H. and Tanimura, M. (1987) ‘The molecular clock runs more slowly in man than in apes and monkeys’, Nature, 326(6108), pp. 93–96. doi: 10.1038/326093a0.

Lim, E. T. et al. (2017) ‘Rates, distribution and implications of postzygotic mosaic mutations in autism spectrum disorder’, 20(9). doi: 10.1038/nn.4598.

Lim, J. and Luderer, U. (2011) ‘Oxidative Damage Increases and Antioxidant Gene Expression Decreases with Aging in the Mouse Ovary’, Biology of Reproduction, 84(4), pp. 775–782. doi: 10.1095/biolreprod.110.088583.

Lindahl, T. and Nyberg, B. (1974) ‘Heat-induced deamination of cytosine residues in deoxyribonucleic acid’, Biochemistry, 13(16), pp. 3405–3410. doi: 10.1021/bi00713a035.

Lindsay, S. J. et al. (2016) ‘Striking differences in patterns of germline mutation between mice and humans’, bioRxiv.

Lister, R. et al. (2009) ‘Human DNA methylomes at base resolution show widespread epigenomic differences’, Nature. Nature Publishing Group, 462(7271), pp. 315–322. doi: 10.1038/nature08514.

Lynch, M. (2010) ‘Evolution of the mutation rate’, Trends in Genetics. Elsevier Ltd, 26(8), pp. 345–352. doi: 10.1016/j.tig.2010.05.003.

Lynch, M. et al. (2016) ‘Genetic drift, selection and the evolution of the mutation rate’, Nature Reviews Genetics. Nature Publishing Group, 17(11), pp. 704–714. doi: 10.1038/nrg.2016.104.

Makova, K. D. and Li, W. H. (2002) ‘Strong male-driven evolution of DNA sequences in humans and apes’, Nature, 416(6881), pp. 624–626. doi: 10.1038/416624a.

Martin, A. P. and Palumbi, S. R. (1993) ‘Body size, metabolic rate, generation time, and the molecular clock.’, Proceedings of the National Academy of Sciences of the United States of America, 90(9), pp. 4087–4091. doi: 10.1073/pnas.90.9.4087.

Mathieson, I. and Reich, D. (2017) ‘Differences in the rare variant spectrum among human populations’, PLoS Genetics, 13(2), pp. 1–17. doi: 10.1371/journal.pgen.1006581.

Mayer, W. et al. (2000) ‘Demethylation of the zygotic paternal genome’, Nature, 403(6769), pp. 501–502. doi: 10.1038/35000656.

Montgomery, S. B. et al. (2013) ‘The origin, evolution, and functional impact of short insertion - deletion variants identified in 179 human genomes’, pp. 749–761. doi: 10.1101/gr.148718.112.Freely.

Moorjani, P. et al. (2016) ‘Variation in the molecular clock of primates’, Proceedings of the National Academy of Sciences, 113(38), pp. 10607–10612. doi: 10.1073/pnas.1600374113.

Moorjani, P., Gao, Z. and Przeworski, M. (2016) ‘Human Germline Mutation and the Erratic Evolutionary Clock’, PLoS Biology, 14(10), pp. 11–13. doi: 10.1371/journal.pbio.2000744.

Müller, H.. (1954) ‘The nature of genetic effects produced by irradiation’, Radiation Biology, (7), pp. 351–473.

Nielsen, C. T. et al. (1986) ‘Onset of the release of spermatozia (supermarche) in boys in relation to age, testicular growth, pubic hair, and height’, Journal of Clinical Endocrinology and Metabolism, 62(3), pp. 532–535. doi: 10.1210/jcem-62-3-532.

Polani, P. E. and Crolla, J. A. (1991) ‘A test of the production line hypothesis of mammalian oogenesis’, pp. 64–70.

Poulos, R. C., Olivier, J. and Wong, J. W. H. (2017) ‘The interaction between cytosine methylation and processes of DNA replication and repair shape the mutational landscape of cancer genomes’, 45(13), pp. 7786–7795. doi: 10.1093/nar/gkx463.

Rahbari, R. et al. (2016) ‘Timing, rates and spectra of human germline mutation’, Nature Genetics. Nature Publishing Group, 48(2), pp. 126–133. doi: 10.1038/ng.3469.

Reik, W., Dean, W. and Walter, J. (2001) ‘Epigenetic reprogramming in mammalian development.’, Science, 293(5532), pp. 1089–1093.

Sasani, T. et al. (2018) ‘Multigenerational CEPH/Utah pedigrees reveal mutation rate variability and quantify post-zygotic and mosaic mutation’, In preparation.

Schultz, M. D. et al. (2015) ‘Human body epigenome maps reveal noncanonical DNA methylation variation’, Nature, 523(7559), pp. 212–216. doi: 10.1038/nature14465.

Ségurel, L., Wyman, M. J. and Przeworski, M. (2014) ‘Determinants of Mutation Rate Variation in the Human Germline’, Annual Review of Genomics and Human Genetics, 15(1), pp. 47–70. doi: 10.1146/annurev-genom-031714-125740.

Seplyarskiy, V. B. et al. (2017) ‘Error-prone bypass of DNA lesions during lagging strand replication is a common source of germline and cancer mutations’, bioRxiv.

Smith, T. B. et al. (2013) ‘The presence of a truncated base excision repair pathway in human spermatozoa that is mediated by OGG1’, Journal of Cell Science, 126(6), pp. 1488–1497. doi: 10.1242/jcs.121657.

Strachan, T. and Read, A. (2010) Human Molecular Genetics. 4th edn. Garland Science.

Titus, S. et al. (2013) ‘Impairment of BRCA1-Related DNA Double-Strand Break Repair Leads to Ovarian Aging in Mice and Humans’, 5(M). doi: 10.1126/scitranslmed.3004925.

Tomkova, M. et al. (2018) ‘DNA Replication and associated repair pathways are involved in the mutagenesis of methylated cytosine’, DNA Repair. Elsevier, 62(November 2017), pp. 1–7. doi: 10.1016/j.dnarep.2017.11.005.

Vogel, F. and Rathenberg, R. (1975) ‘Spontaneous mutation in man’, Advances in human genetics, 5, pp. 223–318.

Wei, H. et al. (2015) ‘Age-Specific Gene Expression Profiles of Rhesus Monkey Ovaries Detected by Microarray Analysis’, BioMed Research International, 2015. doi: 10.1155/2015/625192.

Wilson-Sayres, M. A. et al. (2011) ‘Do variations in substitution rates and male mutation bias correlate with life-history traits? a study of 32 mammalian genomes’, Evolution, 65(10), pp. 2800–2815. doi: 10.1111/j.1558-5646.2011.01337.x.

Wong, W. S. W. et al. (2016) ‘New observations on maternal age effect on germline de novo mutations’, Nature Communications. Nature Publishing Group, 7(May 2015), p. 10486. doi: 10.1038/ncomms10486.

Wu, C. I. and Li, W. H. (1985) ‘Evidence for higher rates of nucleotide substitution in rodents than in man.’, Proceedings of the National Academy of Sciences, 82(6), pp. 1741–1745. doi: 10.1073/pnas.82.6.1741.

Zhang, P. et al. (2009) ‘Transcriptome Profiling of Human Pre-Implantation Development’, 4(11). doi: 10.1371/journal.pone.0007844.

